# GPR161 mechanosensitivity at the primary cilium drives neuronal saltatory migration

**DOI:** 10.1101/2025.03.05.641658

**Authors:** Théo Paillard, Ada Allam, Mohamed Doulazmi, Mathieu Hautefeuille, Coralie Fouquet, Liza Sarde, Julie Stoufflet, Nathalie Spassky, Stéphane Nédélec, Isabelle Dusart, Alain Trembleau, Caillé Isabelle

## Abstract

The saltatory migration of neurons is essential for brain formation. Whether mechanical stimuli regulate this process is unknown. Here we show that the primary cilium acts as a mechanical sensor through GPR161. Using an *ex vivo* neuronal migration model and microfluidic assays, we demonstrate that fluid shear stress induces migration *via* the mechanoreceptor GPR161 at the primary cilium, with its mechanosensitive Helix 8 being essential. We demonstrate that GPR161 activates a recently discovered cAMP/PKA signaling pathway leading to the phosphorylation of NDE1, a dynein complex regulator, and microtubule organization to regulate migration. These findings unveil a dynamic primary cilium-based pathway sensing mechanical stimulus to drive cyclic saltatory neuronal migration during brain development.

## Introduction

During development, vertebrate neurons migrate from their site of birth to their final destination. Neuronal migration defects lead to severe brain malformations associated with neurological or psychiatric disorders (*1–3*). Interestingly, migrating neurons exhibit a saltatory pattern of movements, characterized by alternating phases of somal translocation and pauses (*4*). They display a pattern of successive cellular events including the extension of a leading process in the direction of the migration, a movement of the centrosome within this process (centrokinesis, CK), followed by nuclear translocation (nucleokinesis, NK) and a pause, before the whole process resumes (Fig. 1A) (*4*, *5*). The primary cilium (PC), a microtubule-based organelle emanating from the centrosome was recently shown to regulate the directionality and/or dynamics of migrating neurons (*6–11*). The PC itself displays a cyclic behavior, through externalization during CK, and internalization during pauses (Fig. 1A) (*6*, *10*). The PC also acts as a dynamic and cyclic source of cyclic adenosine 3’, 5’ monophosphate (cAMP) that accumulates at the centrosome during the CK and NK, where it activates Protein Kinase A (PKA) (*11*). Interfering with either cAMP production or PKA centrosomal localization decreases the speed of migration with longer pauses and less frequent NK. Therefore, the PC produces cAMP when it is externalized upon CK, and the resulting PKA activation at the centrosome influences downstream targets that regulate the cytoskeleton (*12*). In recent years, the PC’s ability to deflect in response to mechanical stimuli has raised speculation about its mechanosensory functions. Recent studies provided compelling evidence that the PC plays a mechanical role in the left-right organizer to instruct left-right asymmetry (*13*, *14*). Yet, whether PC-mediated mechanosensitivity plays a role during nervous system development is unknown.

**Fig. 1:**
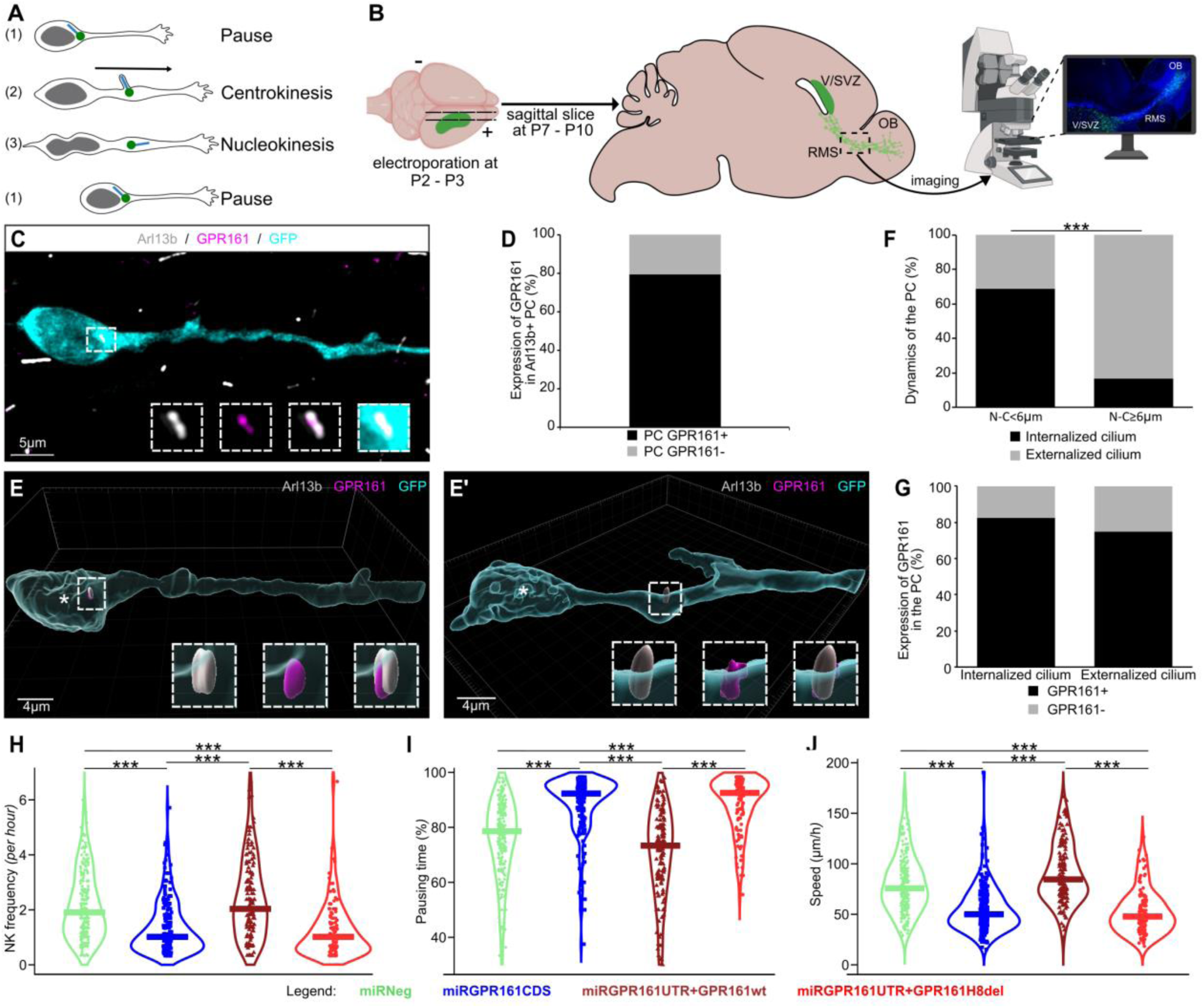
GPR161 regulates migration via its Helix-8. **(A)** Representation of the cyclic saltatory migration steps of a neuroblast. (1) In Pause, the centrosome is next to the nucleus (green) and the primary cilium (PC) is mostly internalized. (2) In centrokinesis, the leading process extends, the centrosome (green dot) moves within a swelling in the leading process, and the PC is externalized at the cell surface. (3) During nucleokinesis, the nucleus moves forward, the PC becomes internalized. **(B)** Experimental procedure: at postnatal day 2-3 (P2-P3), pups were electroporated in the brain ventricles. The brain was taken at P7-P10 to prepare sagittal slices. Last, imaging of the slices was performed. V/SVZ, ventricular/subventricular zone; OB, olfactory bulb; RMS, rostral migratory stream. **(C)** Immunohistochemistry of a GFP-positive RMS neuroblast (cyan) showing GPR161 subcellular immunolabeling (magenta) in the Arl13b-positive PC (gray). Scale bar: 5 μm. **(D)** Percentage of GFP+ RMS-neuroblasts presenting a GPR161+/Arl13b+ or GPR161-/Arl13b+ immunoreactive PC in miRNeg electroporated neuroblasts (N = 3, n = 51). **(E and E’)** Three-dimensional reconstructions of migrating neuroblasts using Imaris software, the immunostaining was performed as in C. The white asterix indicate the center of nucleus. The distance from the nucleus-to-cilium base is less than 6 μm in (E) and greater in (E’). The PC (gray) is GPR161 positive (magenta) and internalized into the cytoplasm in (E) and externalized at the cell surface (E’). **(F)** Percentage of internalized and externalized PC in neuroblasts in a pausing time morphology and a migrating morphology based on the nucleus-to-cilium (N-C) distance. During pausing time (N-C<6μm), the PC is predominantly internalized. During migrating phase (N-C≥6μm), the PC is predominantly externalized at the cell surface (N = 3, n = 53). Pearson’s X2 test (1, N = 53) = 14.50, p <0.001. **(G)** Percentage of Arl13b+/GPR161+ or Arl13b+/GPR161- immunoreactive PCs in their internalized and externalized states (N = 3, n = 51). Pearson’s X2 test (1, N = 51) = 0.432, p = 0.51. **(H-J)** Analysis of the rhythm of migration: **(H)** NK frequency per hour, **(I)** Percentage of neurons in pausing time, and **(J)** speed of migration (µm/hour) in neuroblasts electroporated with miRNeg (green), miRGPR161CDS (dark bue), miRGPR161UTR+GPR161wt (dark red) or miRGPR161UTR+GPR161H8del (bright red). Statistics are presented in fig. S4.

This possibility was supported by the recently described presence of the mechanosensitive orphan Gαs-coupled receptor GPR161 (*15*) in the PC of neurons (*16*). We therefore investigated whether neuronal migration may be mechanically regulated through GPR161-dependent signaling pathways.

## Results

### GPR161 regulates neuronal migration via its mechanosensitive helix 8

The postnatal migration of neuroblasts from the ventricular/subventricular zone (V/SVZ) is a well-established model for studying neuronal migration. In rodents, V/SVZ-derived neuroblasts display a cyclic saltatory migration (Fig. 1A) as they traverse the rostral migratory stream (RMS) to reach the olfactory bulb (OB, Fig. 1B) (*17*). A few days after intraventricular electroporation of a GFP- expressing plasmid in neonate mice, a cohort of migrating neuroblasts can be visualized in acute slices of the RMS through GFP fluorescence (Fig. 1B).

To investigate whether GPR161, a mechano-receptor, has a role in neuronal migration, we first confirmed that GPR161 is expressed by V/SVZ-derived neuroblasts and localized in the PCs *in vivo* (Fig. 1C-D). In GFP+ neuroblasts in the RMS, 79.3% of PC stained by Arl13b, a PC marker (*18*), were also GPR161-immunoreactive (Fig. 1D). Additionally, 3D reconstructions revealed a dynamic PC localization in migrating neuroblasts: it is either internalized (Fig. 1E and <u>movie S1</u>) or externalized (Fig. 1E’ and <u>movie S2</u>). As previously described (*10*), it is internalized during the pausing phases (when the PC is near the nucleus) (69.0%) and externalized during CK (defined here as when the nucleus to primary cilium distance is superior or equal to 6 µm, (*11*)) (83.3%) (Fig. 1F). Interestingly, GPR161 is consistently present in the PC, regardless of its internalization/externalization state (respectively, 82.6% and 75.0%) (Fig. 1G).

We next used an interfering RNA targeting GPR161 mRNA coding sequence (miRGPR161CDS, fig. S1A, fig. S2) to downregulate GPR161 expression in V/SVZ derived neuroblasts. As controls, we used an interfering RNA predicted not to target any known vertebrate mRNA (miRNeg). Five to 7 days after transfection, we imaged the behavior of neuroblasts in acute brain slices of the RMS (miRNeg: <u>movie S3</u>; miRGPR161CDS: <u>movie S4</u>). The parameters of the rhythm of migration (NK frequency, pausing time, and migration speed), were analyzed only for the neuroblasts performing at least 1 NK (i.e. a nuclear movement superior or equal to 6 µm, (*11*)). We detected a significant reduction in NK frequency (expressed as median [interquartile range]: miRNeg: 1.9 [1.8] NK/hour versus miRGPR161CDS: 1.0 [1.1] NK/hour) (Fig. 1H), a significant increase in pausing time (miRNeg: 78.6% [19.3]; miRGPR161CDS: 92.3% [10.2] (Fig. 1I) and a significant decrease in migration speed compared to miRNeg (miRNeg: 75.7 [36.6] μm/hour; miRGPR161CDS: 50.0 [26.9] μm/hour) (Fig. 1J).

These data demonstrate that GPR161 is essential for proper neuronal migration.

GPR161 has been described as mechanosensitive (*15*) and possess a Helix 8 (*19*, *20*), a C-terminal intracellular domain necessary and sufficient for GPCR mechanosensitivity (*21*). To test whether GPR161’s role in migration depends on its mechanosensitivity, we investigated whether the Helix 8 of GPR161 is required for its role in neuronal migration. We created plasmids expressing either the wild-type GPR161 coding sequence (referred to as GPR161wt) or GPR161 lacking its Helix 8 (GPR161H8del) plasmids (fig. S1B-C). To downregulate the endogenous GPR161, but not the exogenous constructs, pups were co-electroporated with GPR161wt or GPR161H8del along with a miRGPR161 targeting specifically the GPR161 3′ untranslated region (miRGPR161UTR) absent from our plasmids sequence (fig. S1B-C). The knockdown efficacy of the miRGPR161UTR was similar to that of the previously used miRGPR161 (fig. S2). Strikingly, the overexpression of the wild-type form of GPR161 in miRGPR161UTR knocked down neuroblasts (miRGPR161UTR + GPR161wt condition, <u>movie S5</u>) rescued all the migration parameters (NK frequency: 2.0 [2.0] NK/hour; pausing time: 73.3% [21.7]; speed: 84.6 [39.0] μm/hour) (Fig. 1H-J). In contrast, the overexpression of the GPR161 lacking the mechanosensitive Helix 8 (miRGPR161UTR + GPR161H8del condition, <u>movie S6</u>) failed to rescue NK frequency, pausing time and speed phenotype induced by the endogenous GPR161CDS knockdown (NK frequency: 1.0 [1.0] NK/hour; pausing time: 92.5% [12.0]; speed: 47.8 [27.0] μm/hour) (Fig. 1H-J). These findings demonstrate that GPR161 is a key regulator of neuronal migration through its Helix 8, suggesting that mechanosensing is implicated in neuronal migration rhythmicity.

### Fluid shear stress induces neuronal migration through GPR161

We then sought to directly test whether mechanical forces influence neuronal migration. As fluid flow-associated shear stress mechanically influences PC dynamics (*22*), we developed a microfluidic device to expose dissociated neuroblasts to controlled fluid shear stress. Neuroblasts isolated from the V/SVZ of neonatal mice (P4-P7) were plated on a 2D Matrigel-coated substrate within a channel connected to pumps that generate a fluid flow (Fig. 2A).

**Fig. 2:**
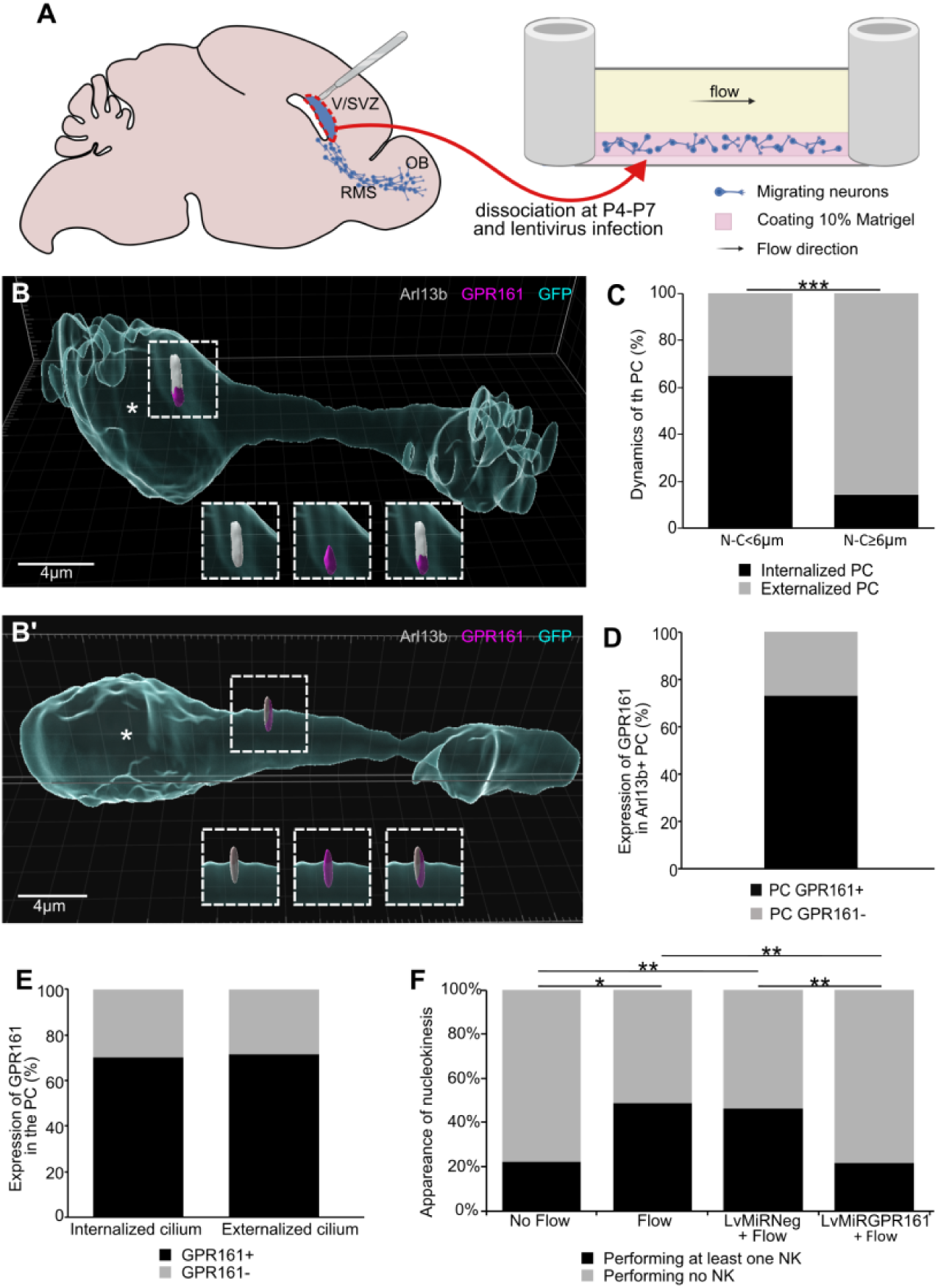
Fluid shear-stress induces migration through GPR161. **(A)** Experimental procedure: The Ventricular/Subventricular Zone (V/SVZ), responsible for generating migrating neuroblasts in the rostral migratory stream (RMS) toward the olfactory bulb (OB), was dissected at P4–P7. Cells were dissociated and infected or not with lentivirus, and then plated in a microfluidic set-up coated with 10% Matrigel. A flow was or not applied. **(B and B’)** Three-dimensional reconstructions of migrating neuroblasts using Imaris software. The white asterix indicate the center of nucleus. The distance from the nucleus-to-cilium base is less than 6 μm in (B) and greater than 6 μm in (B’). The PC (Arl13b positive, gray) is GPR161 positive (magenta) and internalized into the cytoplasm in (B) and externalized at the cell surface in (B’). **(C)** Percentage of internalized and externalized PC in neuroblasts in a pausing time morphology and a migrating morphology based on the distance nucleus-to-cilium (N-C). During pausing time (N-C<6μm), the PC is predominantly internalized. During migrating phase (N-C≥6μm), the PC is predominantly externalized at the cell surface (N = 5, n = 44). Pearson’s X2 test (1, N = 44) = 11.78, p <0.001. **(D)** Percentage of 2D-cultured neuroblasts displaying an Arl13b and GPR161 immunoreactive (GPR161+) or not (GPR161-) PC (N=3, n=26). **(E)** Percentage of Arl13b and GPR161 immunoreactive (GPR161+) or not (GPR161-) PC in its internalized and externalized states (N = 3, n = 24). Pearson’s X2 test (1, N = 24) = 0.006, p = 0.94. **(F)** Percentage of neuroblasts performing at least one nucleokinesis (NK) during the entire movie experiment (2.5 hours) in no-flow condition (N = 3, n = 71), flow condition (N=3, n=75), LvmiRNeg+Flow (N = 3, n = 77) or LvmiRGPR161+Flow (N = 3, n = 65) condition, with a flow corresponding to a shear stress of 0.13Pa. Pearson’s X 2 test (1, N = 142) = 14.30, p = 0.01.

In this set-up, as in brain slices (Fig. 1), neuroblasts had a typical leading process with a PC revealed by the presence of Arl13b immunostaining (fig. S3A and Fig. 2B-B’). 3D reconstructions indicate that neuroblasts PCs retain their typical externalization/internalization dynamics (Fig. 2B and <u>movie S7</u>, and 2B’ and <u>movie S8</u>). As *in vivo* in the postnatal brain (Fig. 1, (*10*)), PCs of neuroblasts in our 2D cultures are internalized during pausing phases in 65.2% of neuroblasts, and externalized during CK in 85.7% of neuroblasts (Fig. 2C). GPR161-immunoreactivity was also detected in 73.1% of Arl13b positive cilium (Fig. 2D), and remained consistently present in the PC, whether it was internalized or not (70.0% GPR161+ in internalized PC and 71.4% GPR161+ in externalized PC (Fig. 2B-B’ and 2E)). Therefore, neuroblasts cultured in 2D in microfluidic chambers retain the subcellular localization of GPR161 and the dynamics of their PC as *ex vivo*. We, therefore, used live imaging in this microfluidic chamber to address the effects of fluid flow-induced mechanical stimulation on neuroblast migration.

Under no-flow conditions, only a small subset of neuroblasts displayed migratory behavior (<u>movie</u> <u>S9</u>), with only 19.7% of the neuroblasts performing one or more NK events over the 2.5-hour time period of imaging (Fig. 2F). In contrast, a fluid flow that generated a 0.13 Pa shear stress doubled the number of migrating neuroblasts, with 40.0% of the cells displaying one or more NK events (<u>movie S10</u>; Fig. 2F). Importantly, analysis of the directionality of NK revealed that this parameter is independent of both the expression of GPR161 and the direction of the flow. Indeed, in the flow conditions, neuroblasts did not migrate more in the flow direction, demonstrating that the migration induced by the fluid flow is not due to a displacement of the neuroblasts by the flow (fig. S3B). Hence, fluid shear stress is sufficient to promote neuronal migration. To determine whether GPR161 is responsible for the fluid flow-dependent activation of migration, we designed a lentivirus co-expressing GFP with either the interfering RNA targeting GPR161 mRNA coding sequence (LvmiRGPR161), or the miRNeg (LvmiRNeg). Upon Lv-MiRNeg infection under flow conditions, neuroblasts displayed the same percentage of cells performing NK as in absence of infection (46.2%, Fig. 2F and <u>movie S11</u>). However, GPR161 knockdown fully blocked the flow induced migration, reverting it to control levels (16.9%; Fig. 2F and <u>movie S12</u>).

Altogether, these results demonstrate that the migration of V/SVZ neuroblasts is regulated by mechanical stimuli and that mechanosensitivity is mediated by GPR161 located at the PC.

### GPR161 regulates the organization of the nuclear cage of microtubules in migrating neuroblasts

We then sought to identify the mechanosensing pathways controlling neuronal migration. As we observed a reduced NK frequency upon GPR161 knockdown (Fig. 1H), we reasoned that GPR161 signaling might regulate cytoskeleton element targets known to be involved in the nuclear movement. In particular, the centrosome, closely linked to the primary cilium, acts as a microtubule organization center extending a microtubule cage around the nucleus, necessary for NK (*23–26*). We thus monitored the organization of the microtubule cage upon GPR161 knockdown. To label the microtubule cage, we co-electroporated with the miRs a plasmid expressing doublecortin (DCX) fused to red fluorescent protein (RFP) (DCX-RFP) (*27*), which was detected by immunofluorescence on fixed brain sections.

Qualitative differences in the organization of the cage of microtubules encircling the nucleus led us to quantify and compare two selected features of this cage in both conditions. First, we distinguished between cages formed exclusively by straight microtubule bundles from those containing bent microtubule bundles (Fig. 3A). Second, we considered the aspect of microtubule bundles at the rear of the nucleus, distinguishing rear-fasciculated vs. rear-disorganized bundles (Fig. 3B). Our quantitative analyses show that while a great majority of miRNeg electroporated neuroblasts display cages with straight bundles of microtubules (71.73% straight, 28.27% bent), a majority of miRGPR161CDS electroporated neuroblasts display bent bundles of microtubules (34.42% straight, 65.58% bent) (Fig. 3C). A similar inversion in the proportion of rear-fasciculated vs. rear-disorganized was observed between miRNeg and miRGPR161CDS electroporated neuroblasts (65.52% rear-fasciculated and 34.48% rear-disorganized in miRNeg; 32.26% rear- fasciculated and 67.74% rear-disorganized in miRGPR161CDS) (Fig. 3D). These results demonstrate that the knockdown of GPR161 in migrating neuroblasts significantly perturbs the organization of their perinuclear cage of microtubules.

**Fig. 3:**
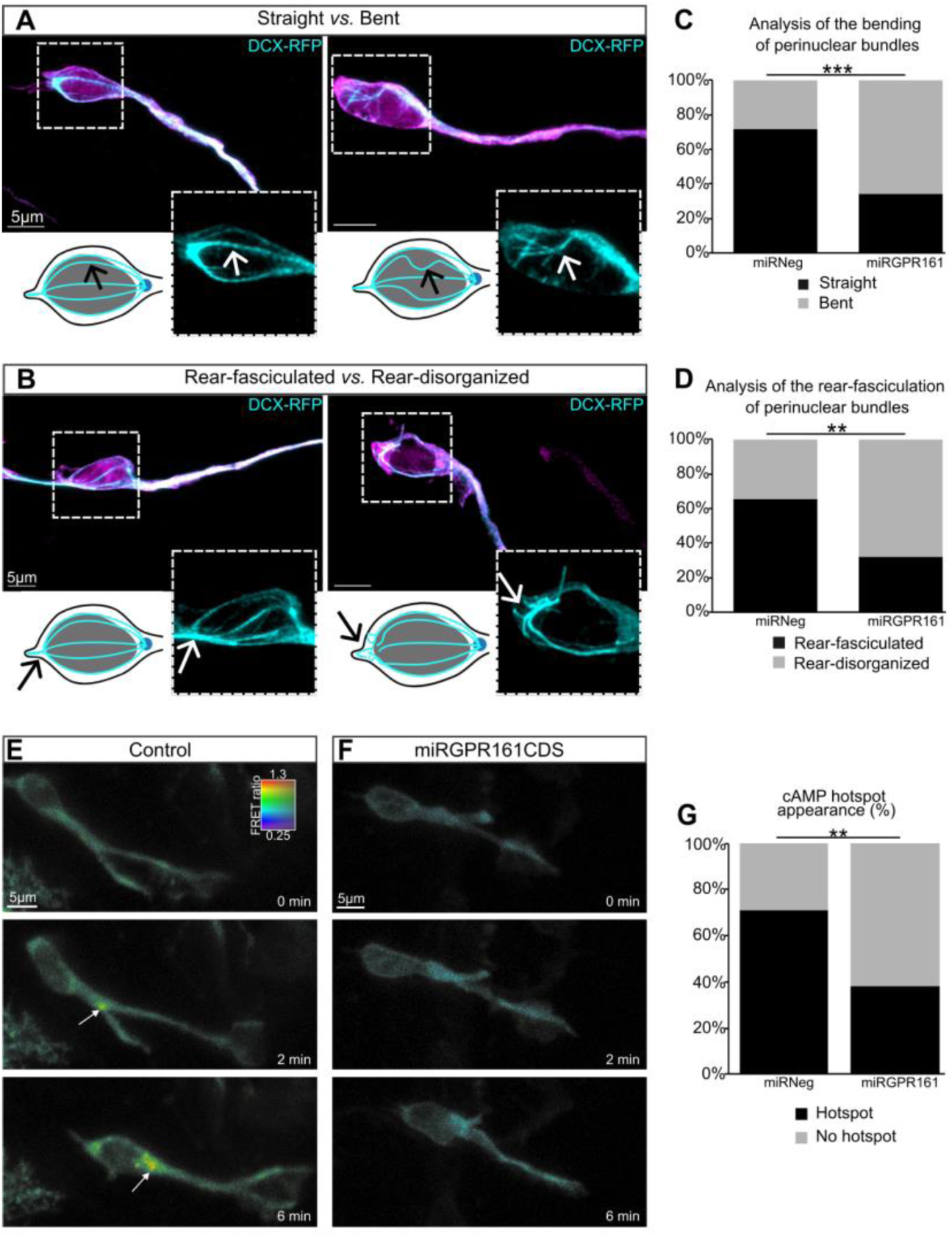
The ciliary receptor GPR161 regulates the organization of the microtubular nuclear cage and is the upstream receptor of cAMP signaling at the centrosome. **(A-B)** Immunohistochemistries of migrating neuroblasts in the RMS electroporated with either miRNeg or miRGPR161CDS (GFP immunolabelling is in magenta) and with DCX-RFP (cyan) labelling the microtubular nuclear cage. Scale bars: 5µm. **(A)** Illustrations of migrating neuroblasts presenting a well-defined straight mictrotubular cage (left, from a miRNeg electroporated neuroblast) compared to a bent microtubular cage (right, from a miRGPR161CDS electroporated neuroblast). (B) Illustrations of migrating neuroblasts presenting a well-defined rear-fasciculated microtubular cage (left, from a miRNeg electroporated neuroblast) compared to a rear-disorganized microtubular cage (right, from a miRGPR161CDS electroporated neuroblast). **(C-D)** Analysis of the percentage of the different microtubular cage defects in miRNeg (N = 3, n = 46) and miRGPR161CDS (N = 5, n = 61) conditions. **(C)** Analysis of the straight vs. bent microtubular cages. Pearson’s X2 test (1, N = 107) = 14.606, p<0.001. **(D)** Analysis of the rear-fasciculated vs. rear-disorganized microtubular cages. Pearson’s X2 test (1, N = 60) = 6.637, p<0.01. **(E-F)** Live two-photon imaging of representative control (E) and miRGPR161CDS (F) neuroblasts co-electroporated with Epac-Sh187 cAMP biosensor. Scale bars: 5 µm. **(E)** White arrow shows a dynamic cAMP hotspot present during NK in the control condition. **(F)** The hotspot is not detected in the miRGPR161CDS condition. **(G)** Percentage of migrating neuroblasts with or without presence of a cAMP hotspot in control (N = 9, n = 52) and miRGPR161CDS conditions (N = 6, n = 34). Pearson’s X2 test with Yate’s continuity correction (1, N = 86) = 7.8509, p = 0.005.

### GPR161 activates the primary cilium-dependent cAMP signaling leading to a cyclic cAMP hotspot at the centrosome

GPR161 is a GPCR coupled to Gs, hence able to trigger the production of cAMP through Gs-dependent activation of an adenylate cyclase (*16*). Interestingly, we recently reported the cyclic appearance of a cAMP hotspot at the centrosome of migrating neuroblasts during CK and NK (*11*). We further showed that ablation of the primary cilium, or knockdown of the ciliary adenylate cyclase 3 (AC3) prevented the production of this cAMP hotspot at the centrosome, and led to migration defects including decreased NK frequency, increased pausing time, and decreased migration speeds (*11*). This migratory phenotype was identical to the one induced by GPR161 knockdown (Fig. 1H-J). This suggested that GPR161 could be upstream of this previously described cyclic accumulation of cAMP.

To test this hypothesis, we co-electroporated the miRGPR161CDS with a FRET cAMP-specific biosensor (*11*) in postnatal mice and imaged cAMP signal in brain slices (Fig. 3F). While 71.2% of control cells displayed a cAMP hotspot (Fig. 3E and <u>movie S13</u>, Fig. 3G), only 38.2 % of miRGPR161CDS displayed a cAMP hotspot (Fig. 3F and <u>movie S14</u>, Fig. 3G). This indicates that GPR161 knockdown abolishes the accumulation of cAMP at the base of the cilium in a majority of migrating neuroblasts. Overall, ciliary GPR161 is the GPCR upstream of the cAMP signaling leading to a cyclic appearance of a cAMP hotspot at the centrosome.

### GPR161 signaling pathway regulates migration through cAMP/PKA-dependent phosphorylation of NDE1

We next ask how the GPR161-dependent centrosomal cAMP accumulation could influence the integrity of the microtubule nuclear cage. One target of cAMP at the centrosome is Protein Kinase A (PKA). We previously showed that delocalization of PKA from the centrosome phenocopies the defects induced by PC deletion or AC3 knockdown (*11*) indicating that centrosomal cAMP/PKA could be key for NK. Interestingly, one centrosomal target of PKA is nudE neurodevelopment protein 1 (NDE1) (*28*). Unphosphorylated NDE1 belongs to a multiprotein complex containing lissencephaly-1 (LIS1) and dynein (*29*), two proteins crucial for the microtubule-dependent nucleokinesis (*30*). This complex plays a key role in neuronal migration (*31–33*).

To determine whether NDE1 may be involved in the GPR161 pathway regulating migration, we first analyzed the localization of NDE1, with a special focus on the centrosome. The high density of chain migrating neuroblasts in the RMS prevented the accurate characterization of NDE1 immunolocalization in brain tissue sections. We thus developed an *ex vivo* culture system, in which V/SVZ explants from postnatal electroporated mice were dissected and cultured in Matrigel for 4 days as previously described (*34*), followed by γ-Tubulin (as a marker of the centrosome) and NDE1 immunodetection (Fig. 4A).

**Fig. 4:**
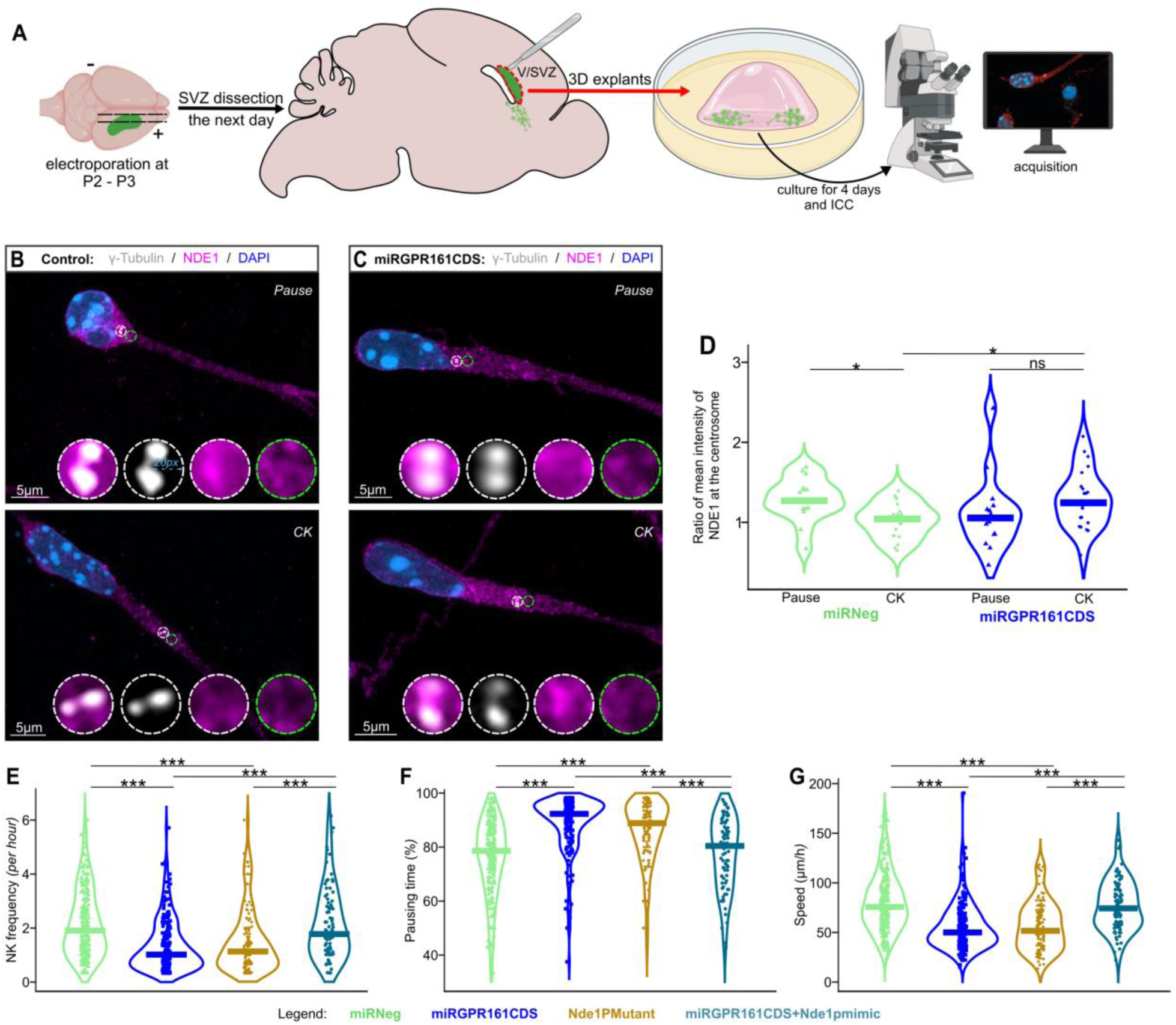
NDE1 is a downstream target of the cAMP/PKA signaling at the centrosome. **(A)** Experimental procedure of 3D explants. V/SVZ, ventricular/subventricular zone; OB, olfactory bulb; RMS, rostral migratory stream, ICC Immunocytochemistry. **(B-C)** Illustrations of a control miRNeg (B) and miRGPR161CDS (C) electroporated neuroblast during pausing time (Pause, top panels) and during centrokinesis phase (CK, bottom panels), showing NDE1 subcellular localization (magenta) and its concentration at the centrosome labelled by γ-Tubulin (gray), nuclear Dapi revelation in blue. The measurement of NDE1 mean intensity at the centrosome was performed within a circle of 20-pixel radius, centered on the centrosome (white circle), and normalized with a same-size circle adjacent in the leading process direction (green circle). **(D)** Analysis of the ratio of the mean intensities of NDE1 at the centrosome level in miRNeg (N = 3) and miRGPR161CDS (N = 4) neuroblasts, during pausing time (Nuclear-Centrosomal distance: N-C<6µm; miRNeg: n = 15, miRGPR161CDS: n = 14) and CK phase (N-C≥6µm; miRNeg: n = 21, miRGPR161CDS: n = 23). Two-way anova: genotype effect (F 1,69 = 1.203, P = 0.276), CK effect (F 1,69 = 0.298, P = 0.586) and genotype CK interaction (F 1,69 = 7.229, P = 0.009) followed by a Benjamini-Hochberg post hoc test. **(E-G)** Analysis of the rhythm of migration: **(E)** NK frequency per hour, **(F)** Percentage of neurons in pausing time, and **(G)** speed of migration (µm/hour), in neublasts electroporated with miRNeg (green), miRGPR161CDS (dark blue), Nde1PMutant (brown) or miRGPR161CDS+Nde1pmimic (light blue). Statistics are presented in fig. S4.

We determined the ratio of NDE1 mean intensity at the γ-Tubulin-labeled centrosome to that in the cytoplasm adjacent to the centrosome (see white and green circles in Fig. 4B and 4C). In control neuroblasts (miRNeg) during the pausing phase (nucleus-to-centrosome distance <6 µm), NDE1 labeling was enriched at the centrosome (most cells had a ratio >1, mean intensity: 1.3 ± 0.3) (Fig. 4B, 4D). However, during CK (nucleus to centrosome distance superior or equal to 6 µm), NDE1 labeling was more homogeneous throughout the cytoplasm, with no enrichment at the centrosome (ratio ∼1, intensity: 1.0 ± 0.3) (Fig. 4C, 4D). This indicates that in control neuroblasts, the centrosomal enrichment of NDE1 during the pause phase is lost during CK (Fig. 4D). In GPR161- knockdown neuroblasts, the NDE1 ratio at the centrosome versus the adjacent cytoplasm was more variable compared to controls and, notably, showed no significant difference between the pause and CK phases (intensity during pause: 1.05 ± 0.4; intensity during CK: 1.24 ± 0.4) (Fig. 4C, 4D). Interestingly, the majority of miRGPR161CDS neuroblasts in CK exhibited centrosomal enrichment of NDE1 compared to control miRNeg neuroblasts in CK (mean ratio 1.27, P<0.05) (Fig. 4C, 4D).

Altogether, these data suggest that activating the GPR161 pathway during migration influences NDE1 localization at the centrosome.

The perturbed centrosomal localization of NDE1 in GPR161 knockdown condition (Fig. 4C-D) prompted us to interrogate whether PKA-dependent phosphorylation of NDE1 is involved in the GPR161-dependent regulation of migration. PKA phosphorylates NDE1 at threonine 131, which decreases its interaction with LIS1 (*28*). To test if NDE1 phosphorylation is involved in neuronal migration, we analyzed in brain slices the effect on neuronal migration of a mutated form of NDE1 which cannot be phosphorylated by PKA (Nde1PMutant, with an alanine substitution of the threonine at position 131, fig. S1D and (*28*)) (Fig. 3E-G and <u>movie S15</u>). Strikingly,

NDE1PMutant induced the same migratory defects as observed upon GPR161 knockdown: reduced NK frequency (1.1 [1.2] NK/hour), increased pausing time (88.9% [14.5]) and reduced migration speed (51.6 [35.4] μm/hour) (Fig. 3E-G). Finally, we tested whether a constitutive phospho-like state of NDE1 could rescue GPR161 knockdown in which NDE1 is delocalized. For that, we co-electroporated the miRGPR161CDS plasmid with a phosphomimetic form of NDE1 (Nde1pmimic, with a glutamate substitution of threonine 131, fig. S1D and (*28*)). In this condition, all the migration parameters were rescued (NK frequency: 1.8 [1.8] NK/hour; pausing time: 80.2% [21.6]; speed: 74.3 [35.4] μm/hour) (Fig. 4E-G and <u>movie S16</u>).

Collectively, these findings indicate that GPR161 and NDE1 phosphorylation at threonine 131 are part of the cAMP/PKA signaling pathway that regulates neuronal migration. Based on these findings, we propose a pathway initiated by the ciliary GPR161, which can act as a mechanosensor, as the primary cilium is externalized at the onset of CK. This activation triggers adenylate cyclase 3 (AC3) activation via Gαs, resulting in the generation of a cAMP hotspot at the centrosome, which leads to NDE1 phosphorylation by PKA, regulating MT nuclear cage organization (Fig. 5).

**Fig. 5:**
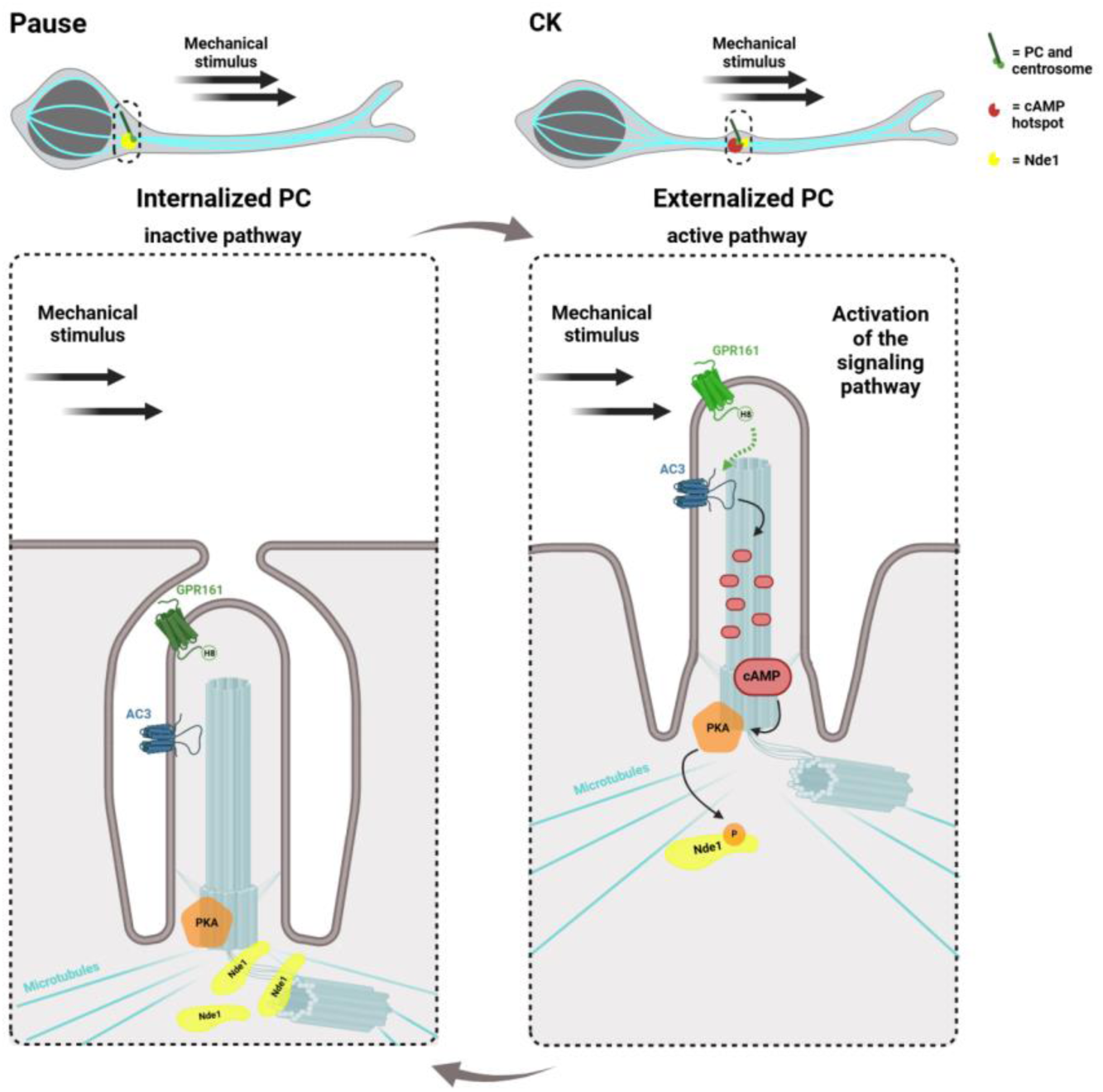
Scheme of cAMP/PKA/NDE1 pathway activation dependent on GPR161’s mechanosensitivity at the primary cilium during centrokinesis. Migrating neuroblasts alternate between phases of pause and centrokinesis (CK). During pausing time, the primary cilium (PC) is internalized and the pathway remains in an inactive state. During centrokinesis (CK), the PC attached to the centrosome moves into a proximal leading process in the direction of migration, and the PC is externalized. The PC externalization allows the mechanosensitive receptor GPR161to be exposed to the surface of the neuroblast and thus to mechanical stimuli. Mechanical activation of GPR161 through its Helix 8 triggers the activation of the ciliary AC3, which in turn produces a cAMP hotspot at the centrosome. This cAMP hotspot activates PKA at the centrosome, enabling the phosphorylation of NDE1 at its threonine 131 residue. Activation of the pathway leads to a reduction in NDE1 localization at the centrosome.

## Discussion

The primary cilium has emerged as a key mechanosensory organelle, capable of detecting and transducing minute mechanical signals essential for key developmental processes like polarization of kidney epithelial cells or the development of left-right asymmetry (*13*, *14*). Our study demonstrates that the mechanoreceptor GPR161 at the primary cilium regulates neuronal migration. Notably, a previous study shows that knocking out GPR161 results in periventricular heterotopia and polymicrogyria (*35*), suggesting that the absence of GPR161 disrupts neuronal migration. We demonstrate here that the mechanosensory property of GPR161 plays a crucial role in neuronal migration. Indeed, fluid shear stress promotes NK in 2D-cultured migrating neuroblasts, and knockdown of GPR161 abolishes this flow-induced migration. Furthermore, our results reveal that the Helix 8 domain of GPR161, linked to GPCR mechanosensitivity (*21*), is critical for its regulatory function on migration, since the GPR161H8del mutant fails to rescue migration defects. The GPR161 mechanosensitivity at the primary cilium thus drives neuronal saltatory migration. *In vitro*, fluid shear stress activates neuronal migration through mechanical forces on GPR161, localized at the primary cilium which cycles between externalization and internalization. *In vivo*, the immediate extracellular environment of the externalized primary cilia in migrating neuroblasts in the adult RMS are the surrounding cells that are used as substrate for their migration, primarily other migrating neuroblasts (*10*). We thus propose that the primary cilium-dependent regulation of migration is driven by frictional forces generated by cell-cell interactions. The dynamic behavior of the cilium likely modulates the mechanical stimulus on it, which in turn would impose a rhythmic saltatory movement of neuroblasts during migration.

Interestingly, our data identify GPR161 as the upstream receptor of the cAMP/PKA-ciliary pathway previously described (*11*), as its knockdown leads to the disappearance of the cAMP hotspot during neuronal migration. GPR161 was initially described as a basal repressor of the Hedgehog pathway via cAMP signaling (*16*), and previous studies have shown an implication of Hedgehog signaling in neuronal migration (*6*, *36*). Nevertheless, our results strongly suggest that the GPR161 function described here does not involve Hedgehog signaling. Indeed, no Shh-expressing cells were detected in the adult RMS (*37*). Furthermore, canonical Hedgehog signaling activation typically delocalizes GPR161 from the cilium (*16*). In contrast, we observed a consistent presence of GPR161 at the PC in migrating neuroblasts. Our findings thus reveal a novel role for GPR161 upstream of the cAMP/PKA signaling pathway, aligning with recent studies that suggest a broader function for GPR161 beyond Hedgehog signaling, notably in promoting tumor cell invasion in triple-negative breast cancer (*20*, *38*, *39*).

Importantly, we also demonstrate that NDE1 is the downstream target of the pathway. First, knockdown of GPR161 disrupts the cAMP-ciliary signaling pathway and abolishes the difference observed under control conditions, where NDE1 is enriched at the centrosome during the pausing phase but not during NK (Fig. 3). Then, NDE1 has been shown to be phosphorylated at the centrosome by PKA on its threonine 131 (T131) (*28*). Noteworthily, our data showed that the overexpression of a non-phosphorylated form of NDE1 on its T131 leads to migration defects similar to those observed in AC3 knockdown, or PKA delocalization from the centrosome (*11*). In contrast, overexpression of a phosphorylated-like form of NDE1 on its T131 rescues all migration parameters in GPR161 knockdown neuroblasts. These findings are in line with prior studies showing that NDE1 loss of function leads to human lissencephaly (*40*, *41*) and schizophrenia (*33*), further supporting the relevance of this protein in proper neuronal migration and development.

Altogether, our findings highlight a pathway initiated by GPR161 at the primary cilium (PC), potentially activated *via* its Helix 8 by a mechanical stimulus from the external environment as the PC emerges at the onset of neuroblast migrating phase. This activation leads to the activation of adenylate cyclase 3 (AC3) *via* Gαs, resulting in the generation of a localized cAMP hotspot at the centrosome. This cAMP signal activates centrosomal PKA, which phosphorylates NDE1 on its T131 (*28*). It is known that NDE1 forms a complex with Lis1 (*31*, *32*) and that the phosphorylation of NDE1 on its T131 by PKA leads to a decreased association with Lis1 (*28*). Our data suggest that upon activation of the pathway, phosphorylated NDE1 may be released from the centrosome. The dissociation of the complex may allow Lis1 to carry out functions in other cellular compartments, such as at the microtubular cage surrounding the nucleus, where it enhances the processivity of Dynein on microtubules (*31*, *32*). Such an hypothesis is consistent with previous studies that identified the Lis1/Dynein complex at the nuclear cage (*42*) and demonstrated that knockdown of Lis1 or dynein disrupts the microtubule network around the nucleus in cortical neuron cultures (*43*).

To conclude, our study identifies GPR161 as a mechanoreceptor at the primary cilium, regulating neuronal migration via its Helix 8 domain. This mechanism is likely general, as GPR161 regulates the cAMP-ciliary pathway previously shown to regulate diverse forms of neuronal migration (*11*). We also establish NDE1 as a critical downstream effector, as it is phosphorylated by PKA at the centrosome. Our findings reveal a cilium-based dynamic pathway that integrates rhythmic mechanosensitivity to drive neuronal saltatory migration. These results open new avenues for undeciphering the role of mechanics and cilia in normal and pathological development.

## Acknowledgments

We thank Sylvie Schneider-Maunoury, Vanessa Ribes, Hynek Wichterle and Arturo Alvarez-Buylla for expert reading, and the members of the IBPS Imaging and Animal facilities of IBPS for their help. Figure 5 was prepared using Biorender.

## Funding

Provide complete funding information, including grant numbers, complete funding agency names, and recipient’s initials. Each funding source should be listed in a separate paragraph.

Agence Nationale pour la Recherche NotifX ANR-20-CE16-0016 (IC)

Agence Nationale pour la Recherche ATOMy ANR-19-CE16-0002 (SN)

Agence Nationale pour la Recherche AetioSpinoids ANR-23-CE16-0026

Sorbonne Université, Paris, France

Centre national pour la Recherche Scientifique (CNRS), France

Institut National de la Santé et de la Recherche Médicale (INSERM), France

## Author contributions

Conceptualization: IC, TP, AA, MH, ID, NS, SN, AT

Methodology: TP, AA, MD, MH, CF, LS, JS

Investigation: TP, AA, MD, CF, LS, JS

Visualization: TP, AA, MD, MH

Funding acquisition: IC, AT

Project administration: IC, AT

Supervision: IC, ID, AT

Writing – original draft: TP, AA, ID, AT

Writing – review & editing: TP, AA, MH, LS, JS, NS, SN, ID, AT

## Data and materials availability

All data are available in the main text or the supplementary materials.

## Supplementary Materials

Materials and Methods

Figs. S1 to S4

Movies S1 to S16

