## Supplementary material for "GPR161 mechanosensitivity at the primary cilium drives neuronal saltatory migration": Methods and Sup Figs

**The PDF file includes:**

Materials and Methods
Figs. S1 to S4

**Other Supplementary Materials for this manuscript include the following:**

Movies S1 to S16

### Materials and Methods

Mice

E14 pregnant C57BL6-J mice were purchased from Janvier labs and housed in a 12-hour light/dark cycle, in cages containing two females. The postnatal mice were housed in the cages with their mother. Animal care was conducted in accordance with standard ethical guidelines [National Institutes of Health (NIH) publication no. 85-23, revised 1985 and European Committee Guide-lines on the Care and Use of Laboratory Animals 86/609/EEC]. The experiments were approved by the ethic committee (Comité d'Ethique en Expérimentation Animale Charles Darwin C2EA-05 and the French Ministère de l'Education Nationale de l'Enseignement Supérieur et de la Recherche, projects APAFIS#1364-2018021915046521 and APAFIS# #45297-2023102517215274). We strictly performed the approved procedures.

Plasmids and viruses

*Plasmids*

Silencing of GPR161 has been performed using BLOCK-iTTM Pol II miR RNAi Expression Vector kits (Invitrogen) and the RNAi Designer (Invitrogen). All the plasmids were used at concentrations between 4 to 8 μg/μl (0.01% Fast green) for postnatal electroporation. When several plasmids were co-injected, the ratio of each was calculated to reach equimolarity in order to transduce cells. The sequence of the single stranded oligonucleotides for miRGPR161CDS are:
top:TGCTGAATACAGCCAAGAGAGTGTGTGTTTTGGCCACTGACTGACACACACTCTTGGCTGTAT;
bottom:CCTGAATACAGCCAAGAGTGTGTGTCAGTCAGTGGCCAAAACACACACTCTCTTGGCTGTA
TTC.
The double stranded oligos were inserted in a pcDNATM6.2-GW/EmGFP-miR. To produce the pcDNATM6.2-GW/Tdtomato-miR, EmGFP was replaced by TdTomato using the Dra1 restriction enzyme. The resulting constructions were sequenced and validated before use.

The sequence of the single stranded oligonucleotides for miRGPR161UTR are:
top:TGCTGTACAGAAGACAACTGAAGTCAGTTTTGGCCACTGACTGACTGACTTCATGTCTTCTGTA;
bottom:CCTGTACAGAAGACATGAAGTCAGTCAGTCAGTGGCCAAAACTGACTTCAGTTGTCTTCTGT
AC.

Constructs of pRP-CMV>mGpr161-wt and pRP-CMV>mGpr161-h8deleted were designed by VectorBuilder. For the pRP-CMV>mGpr161-h8deleted, only the amino acids NKTVRKELLGMC (from n°344 to n°355) representing the Helix 8 were deleted.

pNDE1T131A (Nde1PMutant) and pNDE1T131E (Nde1pmimic) were kindly given by Nicholas Bradshaw (*28*).

pDCX-RFP was a gift from Joseph Gleeson (Addgene plasmid # 32851; http://n2t.net/addgene:32851; RRID:Addgene_32851).

*Viruses*

The pLV-CMV>EGFP-miRGPR161 and pLV-CMV>EGFP-miRNeg viruses were designed by VectorBuilder. VSV-G pseudotyped third-generation lentivirus were used. The same sequences of the single stranded oligos for miRGPR161 and miRNeg were used as in the plasmids above. When used in microfluidic experiments, viruses were added at a MOI of 1 in the resuspension complete medium, before platting the cells.

Postnatal electroporation

Postnatal electroporation was performed at P2, P3 or P4 in C57BL/6J mouse strains. The postnatal mice were anesthetized by hypothermia. Pseudo-stereotaxic injection (from lambda ML: -0.8, A/P:1.1, D/V: 2 at P2; ML: 1.5, A/P:2, D/V: 2.5 at P3 and ML: 1.5, A/P:2, D/V: 2.5 at P4) (using glass micropipette, Drummond Scientific Wiretrol I 50μL, 5-000-1050) was performed and 2 μL of plasmid (between 3 and 10 μg/μL) were injected. Animals were subjected to 5 pulses of 99.99V during 50 ms separated by 950ms using the CUY21 SC Electroporator and 10 mm tweezer electrode (CUY650-10 Nepagene). The animals were placed on 37°C plates to restore their body temperature before returning with their mother. Animals were considered as fully restored when pups were moving naturally, and their skin color returned to pink.

Acute brain slices

Brain slices from mice aged from P6 to P10 were prepared as previously described (11). Briefly, pups were killed by decapitation and the brain was quickly removed from the skull. 250 µm sagittal brain slices were cut with a VT1200S microtome (Leica). Slices were prepared in the ice-cold cutting solution of the following composition: 125 mM NaCl, 0.4 mM $CaCl_2$, 1 mM $MgCl_2$, 1.25 mM $NaH_2PO_4$, 26 mM $NaHCO_3$, 5 mM sodium pyruvate, 20 mM glucose and 1 mM kynurenic acid, saturated with 5% $CO_2$ and 95% $O_2$. Slices were incubated in this solution for 30 min at room temperature and then placed in a recording solution (identical to the solution used for cutting, except that the $Ca^{2+}$ concentration was 2 mM and kynurenic acid was absent) for at least 30 min at 32°C before image acquisition.

Time-lapse video microscopy of migration

To analyze cell migration, images were obtained with an inverted SP5D confocal microscope (Leica) or an upright two-photon microscope Leica SP5 MPII. Images were taken every 3 min for 2-3h using a 40X/1.25 N.A. objective with 1.5 optical zoom on the inverted confocal microscope and with a 25x/0.95 N.A, objective, 1.86x optical zoom. The temperature in the microscope chamber was maintained at 32°C, for postnatal imaging, and brain slices were continuously perfused with a heated recording solution (see above) saturated with 5% $CO_2$ and 95% $O_2$.
Biosensor images were acquired with an upright two-photon microscope Leica SP5 MPII with a 25x/ 0.95 N.A, objective, 4x optical zoom, and GaAsP hybrid detector. The excitation wavelength was set at 850 nm to excite the mTurquoise2 donor. The two emission wavelengths were acquired simultaneously with filters of 479+/-20 nm and 540+/-25 nm. Image stacks with 1µm intervals were taken every 2 minutes for 1h. The presence of TdTomato, indicative of the presence of miRNA, was assessed with a confocal head. The temperature in the microscope chamber was maintained at 32°C and brain slices were continuously perfused with a heated recording solution (see above) saturated with 5% $CO_2$ and 95% O2.

Microfluidic experiments

Pups were sacrificed by decapitation and the brain was quickly removed from the skull. Subventricular zone (SVZ) tissues were dissected in L-15 medium and incubated in 0.25% Trypsin at 37°C for 5 min. Subsequently, cells were dissociated in DNase solution (1mL L-15 + 500µL FBS + 200µL DNase 0.08mg/mL) and counted using a Malassez counting chamber. Cells were resuspended in complete medium.
Commercial Ibidi channels (µ-Slide VI 0.4) were coated with 10% Matrigel diluted in L-15 and incubated for 45 minutes at 37°C in a humidity-controlled incubator. After rinsing with complete medium, 200,000 cells in 65 µL of medium were added to each channel and placed in a 37°C, 5% $CO_2$ incubator for overnight incubation.
Live imaging was performed the next day using a Zeiss inverted 20x/0.95 N.A. objective and 0.6 optical zoom. The temperature in the microscope chamber was maintained at 37°C with 5% $CO_2$.
Flow experiments were conducted using a microfluidic flow system (Flow EZ345, Fluigent) with the pressure controller set to induce a fluid flow of medium inside a commercial microfluidic channel of known dimensions (µ-Slide VI 0.4, Ibidi). The flow rate was set constant with the controller, in the course of 2.5-hour. The pressure pumps were set to apply a shear stress τ of 0.136 Pa, calculated using the following equation: $\tau = \frac{6\eta Q}{h^2 w}$ where η is the medium fluid viscosity (evaluated at $7.10^{-4}$ Pa.s), $Q$ is the flow rate (consistently measured as 1.2 mL/min and controlled by the pressure pumps and limited by the known channel's dimensions)), h is the microchannel height (400 µm) and $w$ its width (3.8 mm).
Control experiments without flow were conducted under the same conditions as above and with live-imaging for 2.5-hour in the same microscopy system but without connecting the microfluidic pump.

Analysis of neuronal migration

Analyses were performed using ImageJ (NIH Image; National Institutes of Health, Bethesda, MD) software and MTrackJ plug-in. The nucleus of each cell was tracked manually on each time frame during the whole movie. For cell migration, calculation of speed, pausing time and nuclear translocation frequency were performed using the x,y,t coordinates of the nucleus of each cell. Cells were excluded from the analysis if they were tracked during less than 30 min or did not perform any nuclear translocation during the whole tracking. A cell was considered as migrating if it performed a distance superior to 6 µm during a 3-minute interval.

More specifically, *speed* was calculated by summing all the distances traveled by one cell and dividing the total distance by the total time, including time slots (3 min interval) when the cell pauses.

The *pausing time* is calculated as the sum of the time slots during which the cell moves less than 6 µm, hence below the cutoff for an NK.

<u>Analysis of biosensor images</u>
The Epac-SH187 cAMP biosensor is composed of a part of Epac protein coupled to a donor and an acceptor fluorophore. This biosensor displays a high ratio change and excellent photostability for measuring live cAMP concentration in the micromolar range. It switches from a high FRET conformation to a lower FRET conformation upon binding of cAMP. The maximum intensity was projected vertically to form a two-dimensional image. Changes in cAMP concentration were analyzed with the plugin FRETratioFx on ImageJ to create a ratio image of non-FRET over FRET fluorescence intensity, which reports biosensor cAMP activation level. The ratio for each pixel is calculated and converted into a hue value.

<u>Immunohistochemistry</u>
P7-P10 mice were deeply anesthetized with 0.1 mL Euthasol (sodium pentobarbital at 40 mg/mL). Intracardiac perfusions with 4% paraformaldehyde (PFA) were performed. Brains were postfixed overnight in 4% PFA. Three rinses were done with PBS 1X (Gibco 1400-067). 70 µm sagittal slices were cut with VT1200S microtome (Leica). Then, slices were placed for one hour in a saturation solution (10% fetal bovine serum; 0,5% Triton-X in PBS). The primary antibodies used in this study were:

- Chicken anti-GFP (Aves, GFP-1020, 1:2,000)
- Rabbit anti-GPR161 (Proteintech, 13398-1-AP, 1:50)
- Mouse anti-Arl13b (NeuroMab, N295B/66, 1:1,000)
- Mouse anti-Ɣ-tubulin (Sigma-Aldrich, T6557, 1:500)
- Rabbit anti-NDE1 (Proteintech, 10233-1-AP, 1:100)
- Rabbit anti-DsRed (Takara, 632496, 1:500)

For GPR161 immunostaining, a pre-treatment of antigen retrieval was done with an incubation of slices in 1 mL of citrate buffer 10 mM for 15 min at 95°C, followed by rinses with PBS 1X. The antibodies were diluted in the saturation solution. Slices were incubated for 48h at 4°C under agitation with the antibodies. Three rinses of twenty minutes were performed with PBS 1X.
The secondary antibodies used were:

- Anti-chicken IgY alexa Fluor 488 (1:1000, Jackson ImmunoResearch: 703-545-155) against anti-GFP.
- Anti-rabbit alexa Fluor 594 (1:1,000, Jackson ImmunoResearch: 711-587-003) against anti-GPR161 and anti-DsRed.
- Anti-Mouse IgG, Fcg Subclass-2A specific alexa Fluor 647 (1:1,000, Jackson ImmunoResearch: 115-605-206) against anti-Arl13b.
- Anti-Mouse IgG, Fcg Subclass-1 specific alexa Fluor 647 (1:1,000, Jackson ImmunoResearch: 115-585-205) against anti-Ɣ-tubulin.

The antibodies were diluted in saturation solution. Slices were incubated for 1h at room temperature under agitation with the secondary antibody solution. Three rinses with PBS 1X were done. Slices were counter-colored with Hoeschst 1:500 and mounted with Mowiol. Acquisitions were performed using a Zeiss LSM 980 upright microscope at high magnification (objective 63X, zoom 3).
To quantify GPR161 knock-down, an Arl13b-positive PC was considered GPR161 negative when it was clearly immuno-negative or indistinguishable from the background at high magnification (objective 63X, zoom 3).
Three-dimensional reconstructions of migrating neuroblasts were performed using Imaris (Carl Zeiss).

Immunostaining on 2D cultured neuroblasts

For the analysis of 2D cultured neuroblasts, such as those used in microfluidic experiments, same dissociation and coating protocol were used as described above (see *Microfluidic experiments* section). Here, neuroblasts were cultured in 12-well removable slide chambers (Ibidi, 81201). After overnight incubation at 37°C, 5% $CO_2$, cells were fixed in 2% paraformaldehyde for 30 min and then rinsed three times with PBS 1x. Primary antibodies were incubated overnight in the saturation solution at 4°C under agitation. Three rinses (20 min each) were performed with PBS 1X. Secondary antibody was incubated for one hour at room temperature under agitation. Three rinses (20 min each) with PBS 1X were done. Two-dimensional cultures were counter-colored with Hoeschst 1:500 and mounted with Mowiol.

Analysis of microtubular cages

To analyze the organization of the microtubular cage in migrating neuroblasts, pDCX-RFP was co-electroporated with either miRNeg-GFP or miRGPR161CDS-GFP at postnatal stages P2–P3. Immunostaining was performed as described in the above Immunohistochemistry section. High-magnification images of electroporated neuroblasts were acquired using a 63× objective with a 3× zoom. Each neuroblast was qualitatively assessed within stacks and categorized based on two criteria:

Analysis of the bending of the microtubular cage: all microtubule bundles are straight and follow the curvature of the nucleus, or at least one bundle deviates from this curvature and appears bent.

Analysis of the rear-fasciculation of microtubular cage: All microtubule bundles reaching the rear of the nucleus fasciculate at the rear of the nucleus, or at least one bundle fails to fasciculate and appears disorganized.

Immunostaining on SVZ explants in Matrigel

For the immunostaining of NDE1, the SVZ of electroporated mice were dissected as described (*34*). SVZ explants were placed on glass-bottom culture dishes (MatTek Corporation; P35G-0-20-C) within 10 mL of 60% Matrigel (Corning; 356237). After Matrigel solidification (15 min at 37°C, 5% CO2), culture medium was added and the dishes were incubated for 4-5 days at 37°C, 5% CO2. For NDE1 immunostaining, SVZ cultures were fixed in 2% paraformaldehyde for 30 min and then rinsed three times with PBS 1x. Primary antibodies anti-NDE1 and anti-Ɣ-tubulin were incubated during four days in the saturation solution at 4°C under agitation. Five rinses (1 hour each) were performed with PBS 1X. The secondary antibodies were incubated for two hours at room temperature under agitation. Three rinses (1 hour each) with PBS 1X were done. Explants were counter-colored with Hoechst 1:500 and mounted with Mowiol.

Image analysis and semi-quantitative analysis of NDE1 immunoreactivity

NDE1 immunostaining was acquired using a Zeiss LSM 980 upright at high magnification (objective 63X, zoom 3). A circle region of interest (ROI, 40 pixel diameter) centered on centrosome (labelled by γ-Tubulin) was manually defined in Fiji. As a reference, a second ROI (40 pixel diameter) was defined next to the centrosomal ROI in the direction of the leading process. Quantification of NDE1 immunofluorescence was performed on the centrosomal z-section, through calculation of the ratio between the centrosomal ROI mean intensity and the reference ROI mean intensity, and the data were exported as CSV files for further analysis.

Statistical analysis

All data manipulation and statistical analyses were performed using R (version 4.3.2, R Foundation for Statistical Computing, Vienna, Austria). Normality of variable distributions was assessed with the Shapiro-Wilk test, while the Levene test was used to evaluate the homogeneity of variances across groups. For variables that failed the Shapiro-Wilk or Levene tests, non-parametric methods were applied, including the one-way Kruskal-Wallis analysis of variance on ranks, followed by the by Dunn's posthoc test with Benjamini-Hochberg p-value correction or Mann-Whitney rank sum tests for pairwise comparisons. Variables that met normality assumptions were analyzed using one-way ANOVA with Benjamini-Hochberg corrections for multiple comparisons or Student's t-test for two-group comparisons. Full statistical analysis of rhythm of migration parameters are shown in Fig; S4.

The orientation of cell migration trajectories, treated as circular variables, was analyzed between groups using circular analysis of variance based on the likelihood ratio test. Categorical variables were compared using

Pearson's $\chi^2$ test or Fisher's exact test. A p-value of $<0.05$ was considered the threshold for statistical significance. Results are expressed as the median [interquartile range (IQR)], with detailed descriptions of statistical tests provided in each figure legend.

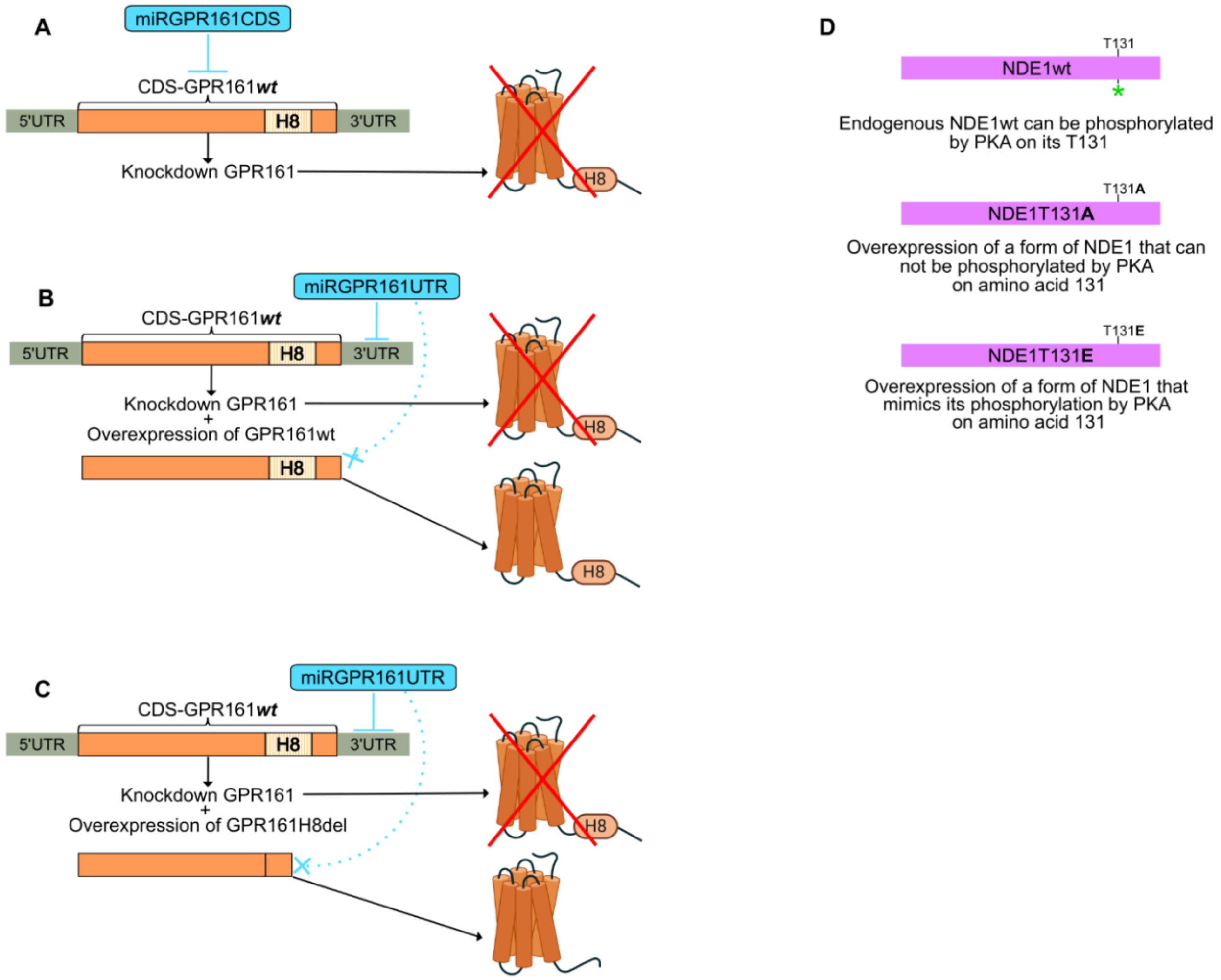


**Fig. S1. Overview of the constructs used in this study.**
**(A)** The miRGPR161CDS plasmid targets the coding sequence (CDS) of endogenous GPR161, leading to its knockdown. **(B)** The miRGPR161UTR plasmid targets the 3′ untranslated region (3′UTR) of endogenous GPR161, also resulting in its knockdown. The plasmid expressing wild-type GPR161 (GPR161wt) is overexpressed and cannot be targeted by miRGPR161UTR due to the absence of a 3′UTR, thereby rescuing GPR161wt expression. **(C)** Similar to (B), the plasmid miRGPR161UTR knocks down endogenous GPR161 by targeting its 3′UTR. Meanwhile, a mutant version of GPR161 lacking Helix 8 (GPR161H8del) is overexpressed. As this mutant also lacks the 3′UTR, it is not targeted by miRGPR161UTR, enabling the expression of the mutant form of GPR161. **(D)** Endogenous NDE1 can be phosphorylated (green asterisk) at the centrosome by PKA on Threonine 131 (T131) (top panel). In the NDE1PMutant condition, we overexpress a non-phosphorylatable NDE1 mutant, where T131 is replaced with alanine (middle panel). In the NDE1PMimic condition, we overexpress a phosphomimetic form of NDE1, where T131 is replaced with glutamic acid to mimic phosphorylation by PKA (bottom panel).

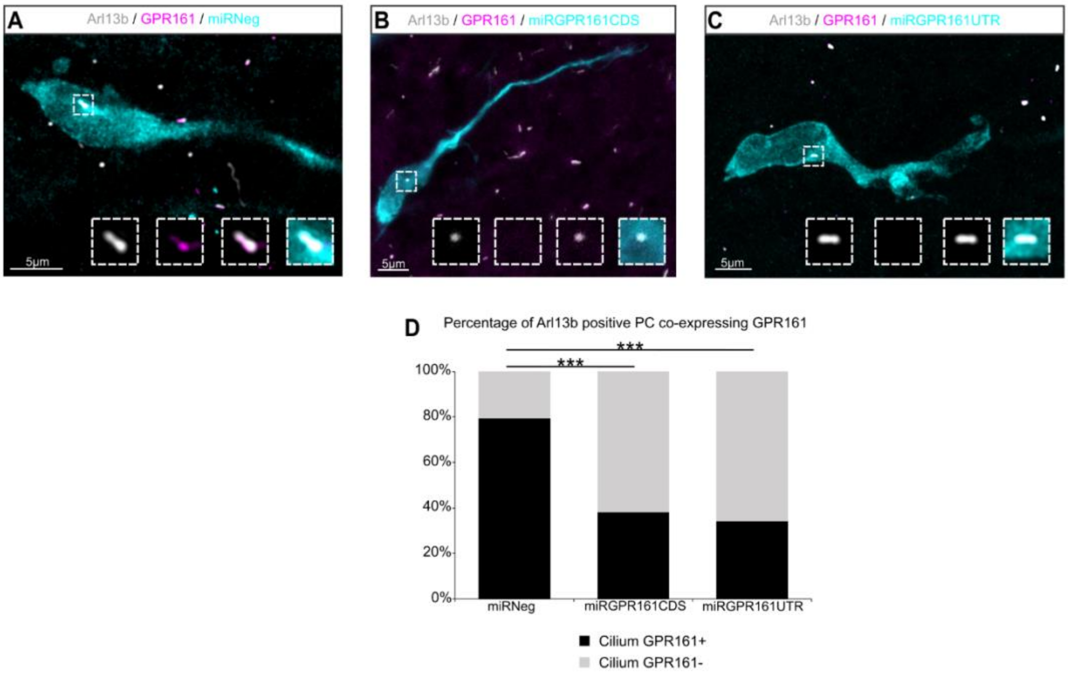


**Fig. S2. Efficiency of miRGPR161CDS and miRGPR161UTR knockdowns.**
**(A-C)** Immunohistochemistry experiments of neuroblasts electroporated with **(A)** miRNeg **(B)** miRGPR161CDSor **(C)** miRGPR161UTR plasmid. **(A)** The miRNeg electroporated neuroblast displays an intact Arl13b immunopositive PC (grey) with GPR161 immunoreactivity (magenta). **(B-C)** Both miRGPR161CDS and miRGPR161UTR display an intact Arl13b immunopositive PC (grey) but without GPR161 immunoreactivity (magenta). Scale bars: 5µm. **(D)** The percentage of electroporated neuroblasts displaying an Arl13b and GPR161 immunoreactive PC is significantly reduced in miRGPR161CDS (38.3%; N=3 n=60) and miRGPR161UTR (34.2%; N=3 n=38) electroporated neuroblasts compared to the control miRNeg (79.3%; N=3 n=53). Pearson's X2 test (2, N = 151) = 25.05, p <0.001.

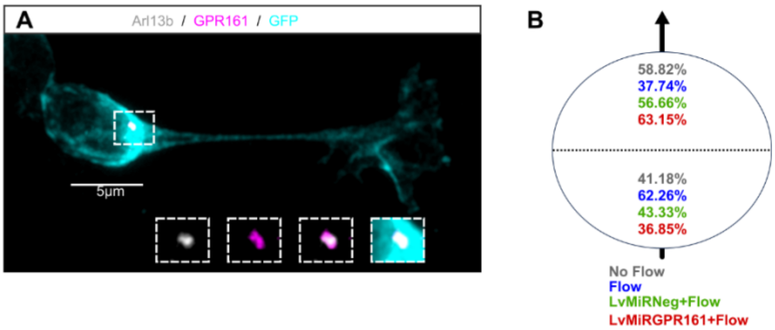


**Fig. S3. Characterization of the 2D culture system and analysis of migration directionality in the microfluidics experiments.**

**(A)** Immunohistochemistry of a 2D-cultured neuroblast (GFP+ cyan) showing GPR161 immunoreactive subcellular expression (magenta) in the Arl13b-positive PC (gray). Scale bar: 5 µm. **(B)** Migration directionality radar represented in 2 spatial dials. Percentage of cells migrating in either spatial direction, relative to the direction of the flow (arrow) in the different conditions: grey, no flow; blue, flow; green, LvMiRneg+flow; red, LvMiRGPR161+flow. Circular analysis of variance based on the likelihood ratio test: $p = 0.60$.

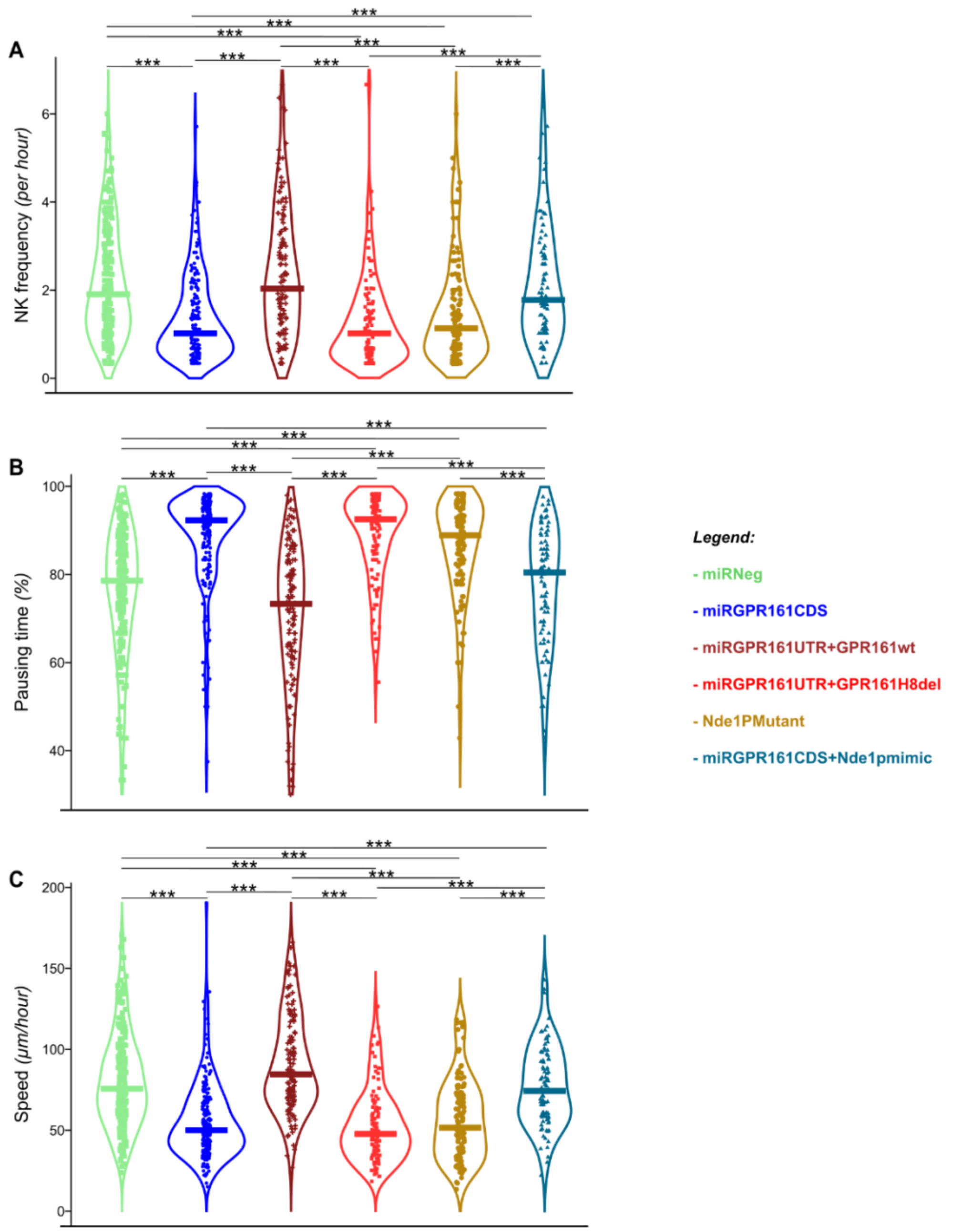


**Fig. S4. Statistical analysis of the rhythm of migration.**

**(A-C)** Migration parameters of neuroblasts electroporated with miRNeg (N = 6, n = 227), miRGPR161CDS (N = 7, n = 203), miRGPR161UTR+GPR161wt (N = 3, n = 139), miRGPR161UTR+GPR161H8del (N = 3, n = 114), Nde1PMutant (N = 4, n = 128) or miRGPR161CDS+Nde1pmimic (N = 3, n = 88) in C57/Bl6 background mice. **(A)** Analysis of NK frequency: miRNeg: 1.9 [1.8] NK/hour; miRGPR161CDS: 1.0 [1.1] NK/hour; miRGPR161UTR+GPR161wt: 2.0 [2.0] NK/hour; miRGPR161UTR+GPR161H8del: 1.0 [1.0] NK/hour; Nde1PMutant: 1.1 [1.2] NK/hour; miRGPR161CDS+Nde1pmimic: 1.8 [1.8] NK/hour. Kruskall-Wallis Test (Chi2 = 130.46, p-value<0.001, df = 2; followed by Dunn's posthoc test with Benjamini-Hochberg p-value correction). **(B)** Analysis of pausing time : miRNeg: 78.6% [19.3%]; miRGPR161CDS: 92.3% [10.2%]; miRGPR161UTR+GPR161wt: 73.3% [21.7%]; miRGPR161UTR+GPR161H8del: 92.5% [12.0%]; Nde1PMutant: 88.9% [14.5%]; miRGPR161CDS+Nde1pmimic: 80.2% [21.6%]. Kruskall-Wallis Test (Chi2 = 242.98, p-value <0.001, df = 2; followed by Dunn's posthoc test with Benjamini-Hochberg p-value correction). **(C)** Analysis of speed of migration miRNeg: 75.7 [36.6] µm/hour; miRGPR161CDS: 50.0 [26.9] µm/hour; miRGPR161UTR+GPR161wt: 84.6 [39.0] µm/hour; miRGPR161UTR+GPR161H8del: 47.8 [27.0] µm/hour; Nde1PMutant: 51.6 [35.4] µm/hour; miRGPR161CDS+Nde1pmimic: 74.3 [35.4] µm/hour. Kruskall-Wallis Test

(Chi2 = 230.84, p-value <0.001, df = 2; followed by Dunn’s posthoc test with Benjamini-Hochberg p-value correction).

**Movie S1.**
Three-dimensional reconstruction image of a neuroblast in the RMS having an internalized primary cilium. Blue, white, and magenta represent cell contour, Arl13b (ciliary marker), and GPR161, respectively. Scale bar: 3µm.

**Movie S2.**
Three-dimensional reconstruction image of a neuroblast in the RMS having an externalized primary cilium. Blue, white, and magenta represent cell contour, Arl13b (ciliary marker), and GPR161, respectively. Scale bar: 3µm.

**Movie S3.**
Time-lapse imaging of miRNeg-GFP electroporated neuroblasts in rostral migratory stream (RMS). The arrow represents the direction of migration, from the V/SVZ to the olfactory bulb. Scale bar: 50 µm.

**Movie S4.**
Time-lapse imaging of miRGPR161CDS-GFP electroporated neuroblasts in rostral migratory stream (RMS). The arrow represents the direction of migration, from the V/SVZ to the olfactory bulb. Scale bar: 50 µm.

**Movie S5.**
Time-lapse imaging of miRGPR161UTR-GFP + GPR161wt electroporated neuroblasts in rostral migratory stream (RMS). The arrow represents the direction of migration, from the V/SVZ to the olfactory bulb. Scale bar: 50 µm.

**Movie S6.**
Time-lapse imaging of miRGPR161UTR-GFP + GPR161H8del electroporated neuroblasts in rostral migratory stream (RMS). The arrow represents the direction of migration, from the V/SVZ to the olfactory bulb. Scale bar: 50 µm.

**Movie S7.**
Three-dimensional reconstruction image of a neuroblast from V/SVZ in 2D culture having an internalized primary cilium. Blue, white, and magenta represent cell contour, Arl13b (ciliary marker), and GPR161, respectively. Scale bar: 3µm.

**Movie S8.**
Three-dimensional reconstruction image of a neuroblast from V/SVZ in 2D culture having an externalized primary cilium. Blue, white, and magenta represent cell contour, Arl13b (ciliary marker), and GPR161, respectively. Scale bar: 3µm.

**Movie S9.**
Time-lapse imaging of 2D-cultured neuroblasts from the V/SVZ under no-flow condition. Tracks highlight neuroblasts with a distinct neuroblast migrating morphology, selected for migration analysis. Scale bar: 50 µm.

**Movie S10.**
Time-lapse imaging of 2D-cultured neuroblasts from the V/SVZ under flow condition (0.13Pa). Tracks highlight neuroblasts with a distinct neuroblast migrating morphology, selected for migration analysis. Flow is directed from the top to the bottom of the field of view. Scale bar: 50 µm.

**Movie S11.**
Time-lapse imaging of 2D-cultured neuroblasts from the V/SVZ infected with LvMiRNeg and under flow condition (0.13Pa). Tracks highlight neuroblasts with a distinct neuroblast migrating morphology, selected for migration analysis. Flow is directed from the top to the bottom of the field of view. Scale bar: 50 µm.

**Movie S12.**

Time-lapse imaging of 2D-cultured neuroblasts from the V/SVZ infected with LvMiRGPR161 and under flow condition (0.13Pa). Tracks highlight neuroblasts with a distinct neuroblast migrating morphology, selected for migration analysis. Flow is directed from the top to the bottom of the field of view. Scale bar: 50 µm.

**Movie S13.**
Time-lapse imaging of a control neuroblast in in rostral migratory stream (RMS) electroporated with EpacSh187 cAMP biosensor. cAMP imaging analyzed with the plugin FRETratioFx on ImageJ to create a ratio image of non-FRET over FRET fluorescence intensity, which reports biosensor cAMP activation level. The ratio for each pixel is calculated and converted into a hue value. Scale bar: 5 µm

**Movie S14.**
Time-lapse imaging of a control neuroblast in in rostral migratory stream (RMS) electroporated with EpacSh187 cAMP biosensor + miRGPR161CDS-Tdto. cAMP imaging was analyzed with the plugin FRETratioFx on ImageJ to create a ratio image of non-FRET over FRET fluorescence intensity, which reports biosensor cAMP activation level. The ratio for each pixel is calculated and converted into a hue value. Scale bar: 5 µm

**Movie S15.**
Time-lapse imaging of Nde1PMutant + miRNeg-GFP electroporated neuroblasts in rostral migratory stream (RMS). The arrow represents the direction of migration, from the V/SVZ to the olfactory bulb. Scale bar: 50 µm.

**Movie S16.**
Time-lapse imaging of Nde1pmimic + miRGPR161CDS-GFP electroporated neuroblasts in rostral migratory stream (RMS). The arrow represents the direction of migration, from the V/SVZ to the olfactory bulb. Scale bar: 50 µm.
